# Uncovering Latent Structure in Gliomas Using Multi-Omics Factor Analysis

**DOI:** 10.64898/2026.03.02.709031

**Authors:** Catarina Gameiro Carvalho, Alexandra M. Carvalho, Susana Vinga

## Abstract

**Background:** Gliomas are the most common malignant brain tumors in adults, characterized by a poor prognosis. Although the current World Health Organization (WHO) classification provides clear guidelines for classifying oligodendroglioma, astrocytoma, and glioblastoma patients, significant heterogeneity persists within each class, limiting the effectiveness of current treatment strategies. With the increase of large-scale multi-omics datasets due to advancements in sequencing technologies, and online databases that provide them, such as The Cancer Genome Atlas (TCGA), it is now possible to investigate these tumors at multiple molecular levels.

**Methods:** In this work, we apply integrative multi-omics analysis to explore the interplay between genomic (mutations), epigenomic (DNA methylation), and transcriptomic (mRNA and miRNA) layers. Our approach relies on Multi-Omics Factor Analysis (MOFA), a Bayesian latent factor analysis model designed to capture sources of variation across different omics types.

**Results:** Our results highlight distinct molecular profiles across the three glioma types and identify potential relationships between methylation and genetic expression. In particular, we uncover novel candidate biomarkers with prognostic value, as well as a transcriptional profile associated with neural system development.

**Conclusions:** These findings may contribute to more personalized therapeutic strategies, potentially enhancing treatment effectiveness and improving survival outcomes for this disease.

## 1 Introduction

Advancements in sequencing technologies have significantly expanded our understanding of biological systems, leading to the generation of vast amounts of high-dimensional data [1]. Various online platforms now provide access to omics data from different diseases, offering valuable resources for biomedical research [1].

Effectively analyzing all of this information requires computational methods capable of handling large datasets. Additionally, to further enhance comprehension, integrating multiple layers of biological data has become crucial, as it leverages interconnected information inherent in different biological domains [2]. Many multi-omics integration methods, with different underlying approaches, have been proposed and validated [3, 4, 5, 6], demonstrating the ability to uncover novel biomarkers and identify molecular signatures, which can then be verified in further molecular researches [7], potentially offering new perspectives for therapeutic intervention.

This is particularly critical in cancer research, which is defined by its inherent heterogeneity and the diverse responses to treatment observed across patients. Gliomas, in particular, represent a challenging class of tumors due to their high degree of molecular heterogeneity [8]. In 2021, the World Health Organization (WHO) redefined the classification of gliomas, due to better reflect advances in molecular biology, allowing a possible early diagnosis [9].

Despite significant advancements in understanding gliomas at the molecular level, many aspects of their biology remain poorly understood, as evidenced by the poor survival rates associated with these tumors.

The existing knowledge gap highlights the need for further investigation into the molecular mechanisms underlying glioma pathogenesis. A deeper understanding of these complexities is essential for the development of more effective, personalized treatment strategies tailored to the unique molecular characteristics of individual patients [10].

## 2 Background

Gliomas are a heterogeneous group of brain tumors, representing one of the most common forms of brain cancer and associated with poor survival outcomes [11]. Over the years, extensive research has aimed to improve diagnosis, prognosis, and treatment strategies for gliomas. Many classifications have been proposed and are continually revised. The most recent classification happened in 2021 by WHO [12], which introduced a revised system incorporating molecular features in addition to histological features, enabling a more accurate and clinically relevant stratification of glioma subtypes, which is crucial for the development of effective targeted therapies [10].

Adult gliomas are classified into three subtypes: *astrocytoma* (grades II, III, IV), *oligodendroglioma* (grades II, III), and *glioblastoma* (grade IV) [9]. The term Lower-Grade Gliomas (LGGs) typically refers to the first two, highlighting their lower aggressiveness compared to glioblastoma (GBM). The mutations in the genes *IDH1* or *IDH2*, present in astrocytomas and oligodendrogliomas, are in fact associated with a better prognosis [9]. What distinguishes astrocytoma from oligodendroglioma, within LGGs, is the presence of the 1p/19q codeletion in oligodendroglioma patients. On the other hand, GBM is a *IDH*-wildtype tumor and is characterized by a gain of chromosome 7 and the loss of chromosome 10 (+7/-10), with other key molecular alterations, including *EGFR* amplification and/or *TERT* promoter mutations. The presence of morphological features such as necrosis, microvascular proliferation, and mitotic activity is also responsible for a higher grade classification [9].

Despite this strict categorization, gliomas exhibit a wide range of molecular profiles [9]. In this context, multi-omics integration analysis plays a crucial role, as it enables a comprehensive view of the tumor biology by combining information from various molecular omics layers, such as genomics, transcriptomics, epigenomics, and proteomics [1, 2]. Its main objectives include discovering molecular mechanisms, clustering samples, and predicting outcomes, such as patient survival or treatment efficacy [7].

To achieve these goals, a variety of multi-omics integration methods have been developed [13]. An important group of strategies can be classified as joint Dimensionality Reduction (jDR) methods that decompose each omics dataset into the product of a factors matrix (shared by all omics) and a loadings matrix unique for each omics dataset [3]. It encompasses correlation and covariance-based approaches aimed at maximizing the covariance or correlation between datasets, such as Canonical Correlation Analysis (CCA) [14] and its variants, as well as Factor Analysis (FA), which assumes latent variables capture the shared variance across the data. Factor analysis can further include Probabilistic/Bayesian Models (PR), such as Multi-Omics Factor Analysis (MOFA) [15] or iCluster [7]. Other approaches include Similarity (Kernel)-based (KB) methods, where a similarity matrix is constructed from kernels and analysed afterwards, and Network-based Integration (NB) methods [16], which represent data as networks, where the nodes are entities and the edges are similarities and interactions. This last approach typically incorporate molecular interaction networks or define custom networks based on omics data. Even though that NB methods generally perform better due to their reliance on a priori biological knowledge [6], which helps reduce false discoveries, this data is often incomplete or unavailable. Deep Learning (DL) models have gained significant attention in recent years, due to their ability to effectively handle high-dimensional data and capture complex non-linear relationships [5]. Even though they have demonstrated superior accuracy compared to traditional methods in downstream tasks such as classification and regression [17], their lack of interpretability remains a significant limitation, despite the integration of explainability techniques into some models [18, 19].

## 3 Materials and Methods

### 3.1 Data Availability

The integration in the present study focused on several omics layers, specifically genomics (mutations), as they are key drivers of glioma heterogeneity, along with transcriptomics (mRNA and miRNA) and epigenomics (DNA methylation). The data was obtained from The Cancer Genome Atlas (TCGA) under the project names ‘TCGA-GBM’ and ‘TCGA-LGG’.

The mutations dataset was downloaded using the `RTCGAToolbox` [20] package, while the others were obtained using the `TCGAbiolinks` [21] package. The downloaded data included binary mutation profiles, count-based transcriptomics, and beta values representing the methylation proportion of each probe. These last two datasets were provided in the *summarizedExperiment* format, containing not only the expression matrix but also features metadata, including gene annotations and additional biological information such as the associated chromosomal locations and the corresponding gene names.

Clinical data, including patient demographics (age, sex) and survival information, were also extracted. Its summary is in Table 1.

**Table 1:** Clinical characteristics of patients in the “TCGA-GBM” and “TCGA-LGG” cohorts. Values are also shown for the full dataset.

|  |  | TCGA-GBM<br>(n= 617) | TCGA-LGG<br>(n = 516) | Total<br>(n = 1133) |
| --- | --- | --- | --- | --- |
| Age, mean (StD) | | 58 (14.41) | 43 ±13.36 | 51 ± 15.78 |
| Sex, n (%) |  |  |  |  |
|  | Female | 230 (37.3) | 230 (44.6) | 460 (40.6) |
|  | Male | 366 (59.3) | 285 (55.2) | 651 (57.5) |
| Vital Status, n (%) |  |  |  |  |
|  | Alive | 101 (16.4) | 388 (75.2) | 489 (43.2) |
|  | Dead | 492 (79.8) | 126 (24.2) | 618 (54.5) |
| Primary Diagnosis, n (%) |  |  |  |  |
|  | Astrocytoma | - | 194 (37.6) | 194 (17.1) |
|  | Oligodendroglioma | - | 190 (36.8) | 190 (16.8) |
|  | Mixed Glioma | - | 131 (25.4) | 131 (11.6) |
|  | Glioblastoma | 599 (97) | - | 599 (52.3) |

The ground-truth gliomas labels used were from the study [22], where the TCGA labels were updated according to the most recent WHO guidelines from 2021. The labels assigned were: ‘Astrocytoma’, ‘Glioblastoma’, ‘Oligodendroglioma’ or ‘Unclassified’.

All the code is available at https://github.com/CatarinaGameiroC/Article-MOFAinGlioma to ensure reproducibility and modularity.

### 3.2 Data Preprocessing

To address missing values in the DNA methylation dataset, features with 90% or more missing entries were excluded. Outliers and low-abundance features were filtered out across datasets to ensure data quality. Each dataset was transformed into continuous values, except for the mutations dataset, which remained binary. Specifically, in the epigenomics data, beta values were converted to M-values, which correspond to the methylation signal, while transcriptomics data were normalized, variance-stabilized, and transformed into logarithmic counts per million (log-CPM). To reduce computational complexity and eliminate redundant or noisy features, low-variance features were filtered: only the top 2% most variable probes were retained for DNA methylation, the top 50% for mRNA, and the top 80% for miRNA.

Lastly, the samples from the four omics datasets were intersected prior to model application, resulting in a final cohort of 318 patients. Among them, 142 are classified as astrocytoma, 80 as glioblastoma, and 84 as oligodendroglioma, with 12 samples left unclassified. Sex and age information was not available for only one patient. Regarding survival data, it is completed for 315 patients.

### 3.3 Multi-Omics Factor Analysis (MOFA)

MOFA is a factor analysis method that operates within a probabilistic Bayesian framework [15]. Given *M* omics datasets represented by *X*^*m*^ with dimension *n × p*_*m*_, where *m* = 1, 2, …, *M, n* is the number of observations and *p*_*m*_ is the number of features in the *m*-th omics dataset. The goal of MOFA is to factorize each matrix into a shared set of *K* latent factors:

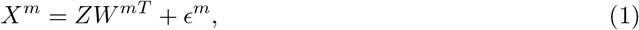

where *Z*_*n×K*_ is called the factors matrix, shared across all of the omics, capturing the low-dimensional latent variables, while 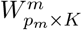 is the loadings matrix that relates the high-dimensional space to the low-dimensional representation. The residual noise, *ϵ*^*m*^, captures variability not explained by the model and allows for heteroscedasticity across features, with each feature having variance 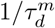.

The prior for *Z* follows a standard normal distribution, ensuring that the latent factors are normally distributed. For the loadings *W* ^*m*^, MOFA incorporates two types of sparsity constraints: one on the factors (View- and factor-wise sparsity) and another on the weights (Feature-wise sparsity). The latter is based on the assumption that biological sources of variability are typically sparse, meaning that only a small number of features are active. This results in the prior given by:

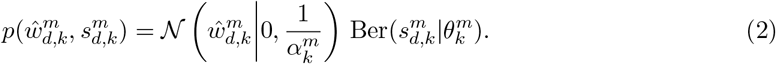

The term 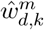 represents the continuous weight component and 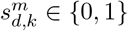 indicates if the weight is active.

Inference is performed using variational inference, where the posterior distribution is approximated through optimization by maximizing the Evidence Lower Bound (ELBO):

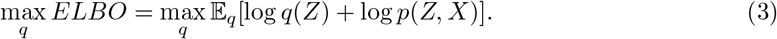

In variational inference, the distribution *q*(*Z*) serves as an approximation of the true posterior *p*(*Z*|*X, θ*). The mean-field approximation further assumes that the variational distribution *q*(*Z*) factorizes over disjoint groups of variables, simplifying the optimization process. In MOFA, this translates into:

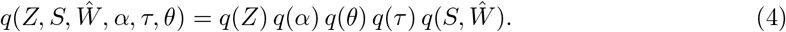

MOFA offers great flexibility by supporting different priors on the residual noise, Gaussian, Bernoulli, or Poisson, allowing it to handle continuous, binary, and count data. However, binary and count data are often not well modeled by these distributions. Count data is usually very sparse, making it preferable to apply data transformations to approximate a Gaussian distribution. Additionally, MOFA naturally accommodates missing values by excluding them from the likelihood computation, thus not affecting the update equations.

MOFA was applied in this work using the R package `MOFA2`. To ensure robustness in MOFA model selection, 10 independent runs were performed, each initialized with a different random seed. The model was configured with sparsity constraints enabled, and the number of factors was not predefined; instead, factors explaining less than 5% of the variance were automatically discarded. The likelihoods were assigned based on data characteristics, with a Gaussian distribution applied to most omics data, while binary likelihoods were used for genomic features. The convergence mode was set to “medium”, stopping when the delta ELBO change reached 0.00005%. After training, the best model was selected using ELBO-based optimization.

### 3.4 Differential Gene Expression Analysis (DGE)

DGE analysis was performed to identify the most significant genes distinguishing glioma subtypes in mRNA and DNA methylation data. For mRNA, the `edgeR` package [23] was used with a treshold of 0.05 for the False Discovery Rate (FDR) and regarding Fold-Change (FC), | log_2_ FC| *>* 1. For methylation, `CHAMP` was applied with FDR *<* 0.05 and a change in mean beta values (Δ*β*) greater than 0.3.

### 3.5 Gene Set Enrichment Analysis (GSEA)

GSEA was performed on ranked gene features from MOFA, using the `ReactomePA` [24] package for mRNA. Top-level pathway information, from data available on the Reactome website, was integrated to enhance biological interpretability. However, since this reference file did not include all top-level associations, missing entries were retrieved using an R interface to the Reactome API [25].

The `missMethyl` package [26] was used for DNA methylation analysis to correct for bias caused by varying CpG coverage across genes. The enrichment focused on Gene Ontology (GO) pathways, focusing on Biological Processes (BP). Only pathways containing between 5 and 500 genes were considered for testing, using a significance threshold of *p <* 0.05, with multiple testing correction applied via the Benjamini–Hochberg (BH) procedure. To focus the analysis on regions with potential regulatory impact, it was also restricted the input to CpGs located within promoter-associated regions, specifically those annotated as *TSS1500, TSS200*, or *1stExon*.

### 3.6 Survival Analyis

To assess statistical differences between two Kaplan–Meier survival curves, the log-rank test was performed with groups defined by median gene expression. This analysis was conducted using the `survival` package [27].

## 4 Results

### 4.1 Overview of the Model

Applying MOFA resulted in the identification of four factors. The first factor accounts for 80% of the total variance, capturing co-variation across all datasets, suggesting its strong relevance. Factor 2 primarily explains variance within the mRNA assay, while factor 3 captures variation across all datasets except for miRNA. Notably, the miRNA assay is poorly represented in the model, contributing only 5.17% of its total variance (Fig. 1).

**Figure 1:**
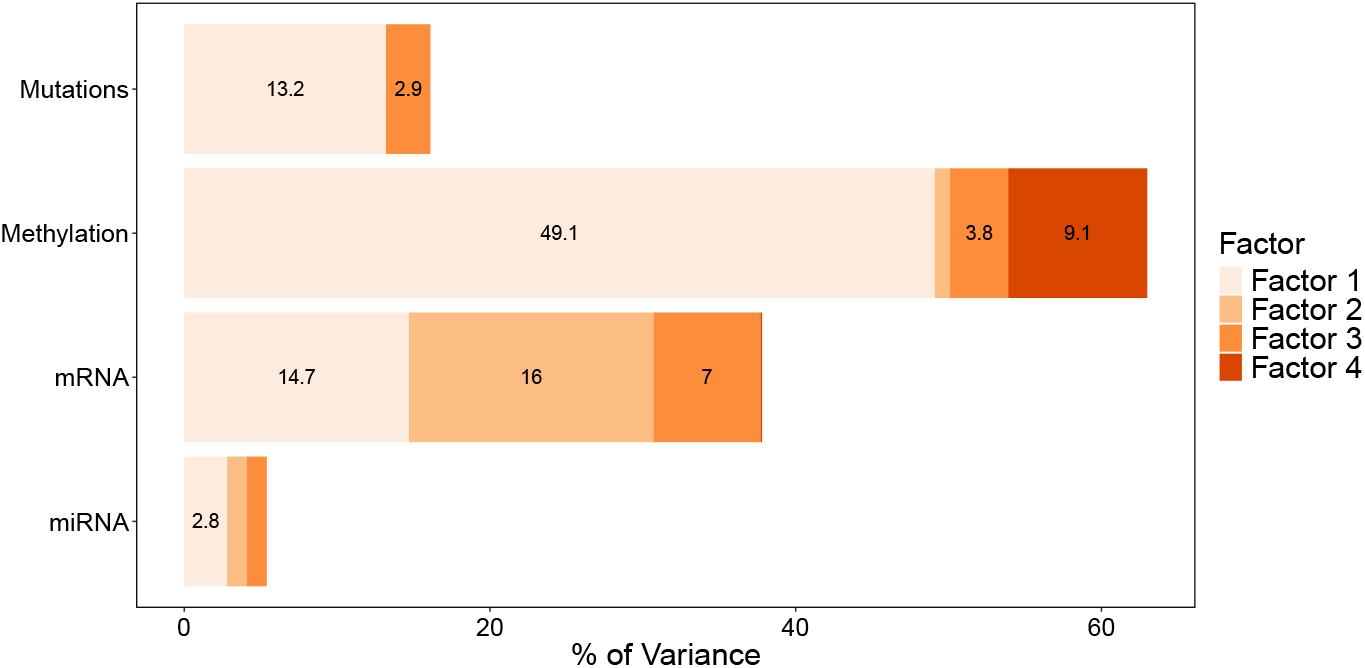
Variance decomposition by omics data type and latent factors. Only variance values for factors explaining more than 2% of the variance within an omics dataset are shown.

Consequently, factors 1 and 3 appear to capture relevant general information of gliomas, while factor 2 reflects a specific gene expression pattern. These three were selected for further analysis, as factor 4 only separates samples by sex. Sample projections of the relevant factors are shown in Fig. 2.

**Figure 2:**
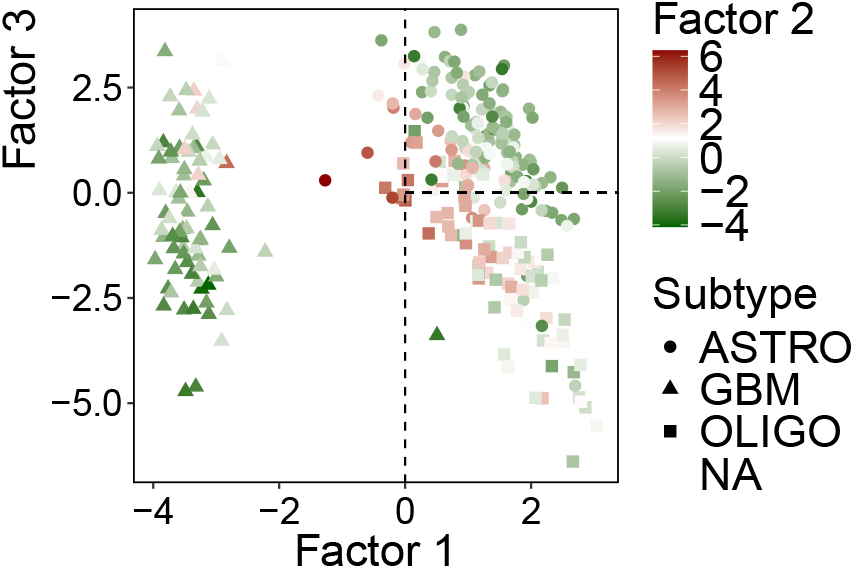
Latent space projections of samples across MOFA factors 1 and 3, shaped by subtype and colored by factor 2 values. Dashed lines indicate the vertical line at Factor 1 = 0 and the horizontal line at Factor 2 = 0 (extending from Factor 1 = 0 onwards).

Fitting a univariate Cox proportional hazards model to each factor individually revealed that the first three factors were significantly associated with survival at a significance level of 0.05. However, when all factors were included simultaneously in a multivariate model, only the first factor remained statistically significant. Univariately, the first factor shows a hazard ratio of approximately 0.52, indicating that a one-unit increase in this factor is associated with a 48% reduction in the hazard of death. This result is consistent with the known aggressiveness of GBM, suggesting that this factor may capture relevant biological signals related to poor prognosis. The results are presented in Fig. 3.

**Figure 3:**
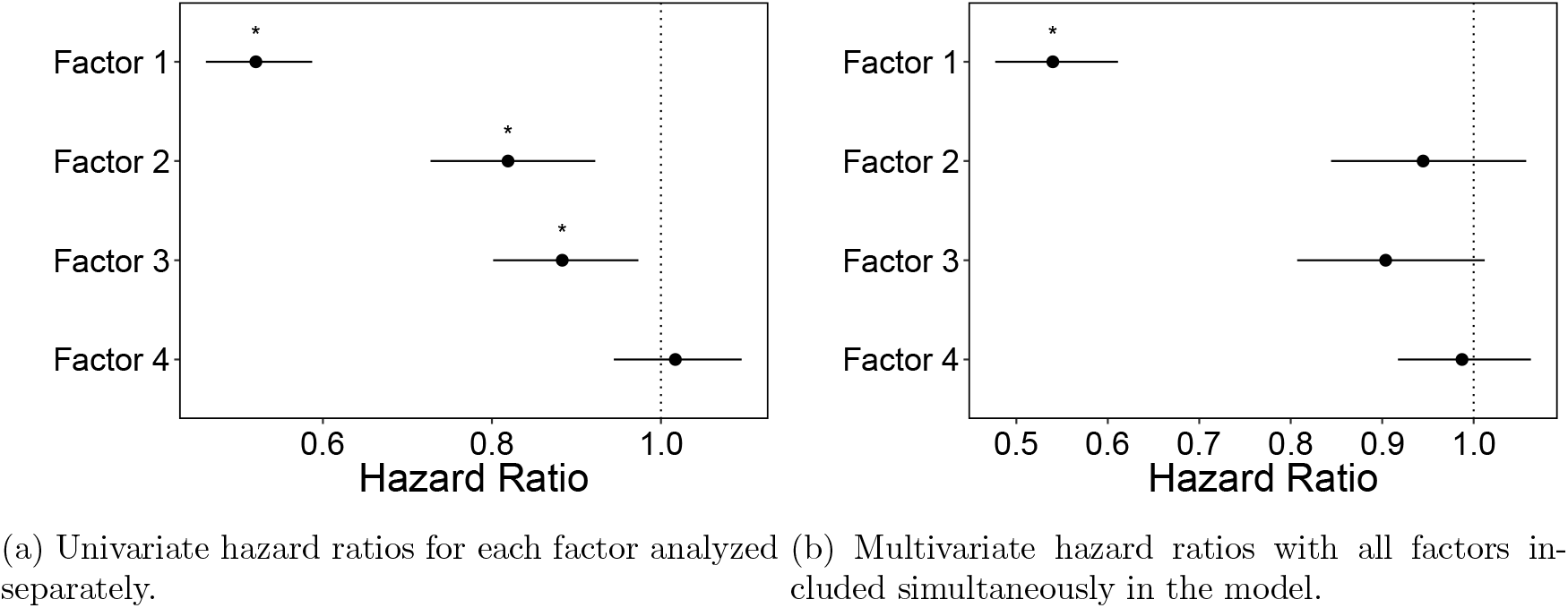
Hazard ratios and corresponding 95% confidence intervals for the four MOFA-derived factors. Statistically significant associations (*p <* 0.05) are marked with an asterisk.

#### 4.1.1 Features Selected

The loadings matrix for mutations and miRNA is notably sparse. It is observed that, for the mutations loadings matrix, factor 1 is strongly associated with gene *IDH1* on the positive side, while it selects *PTEN* and *EGFR* negatively. Factor 3 highlights *ATRX* and *TP53*, while it shows a negative association with *CIC*. Fig. 4 shows the clear separation of mutation profiles by these two factors.

**Figure 4:**
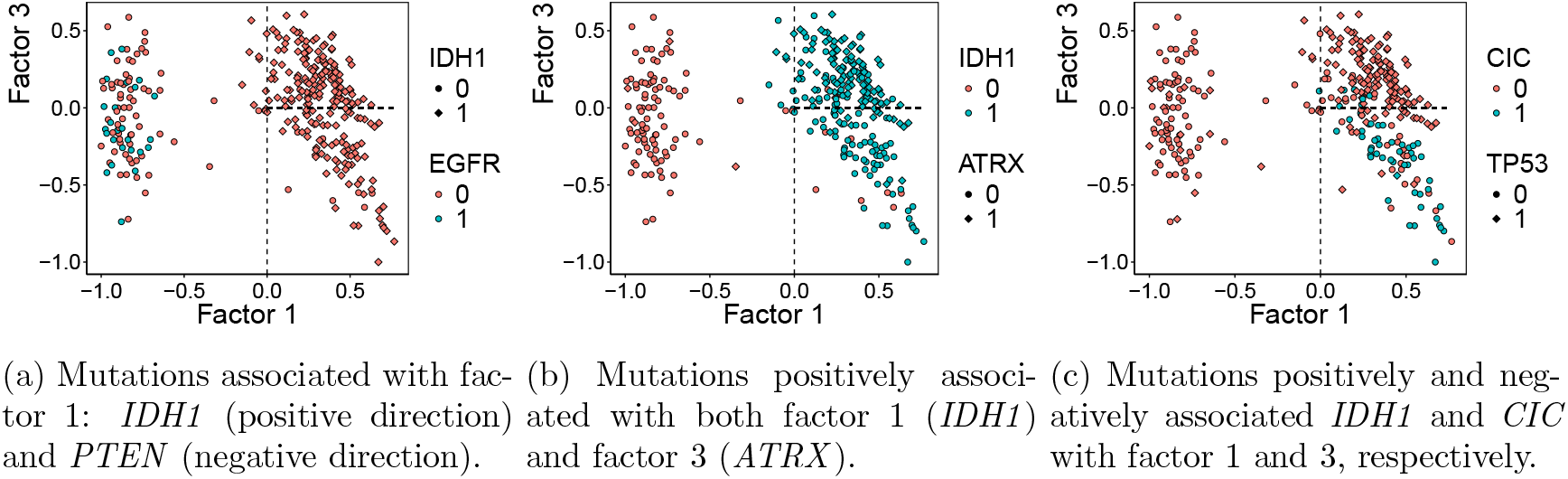
Projection of samples on factors 1 and 3, colored and shaped by driver mutation status.

Considering miRNA, for factor 1 (only expressed in this factor), the following genes *ENSG00000278783, ENSG00000207575, ENSG00000266174* were selected, corresponding to the genes *MIR6071, MIR649* and MIR4666A, respectively, associated with a positive loading.

The loading matrices for mRNA and DNA methylation were not particularly sparse. Therefore, for each of the three factors (1, 2, and 3), the top 30 features with the highest absolute loadings were selected to facilitate a more focused and interpretable analysis. The absolute value of the loadings was considered because, in contrast to mutations, both high and low values in these omics layers (mRNA and DNA methylation) can carry meaningful biological information. Despite this, the selected features generally exhibited consistent sign patterns across factors. All methylation features selected for factor 1 had positive loadings, whereas those for factor 3 had negative loadings. Interestingly, eight probes selected by factor 3 are assigned to the gene *ISM1*. For mRNA, the selected features in factor 1 were mostly negative (except for three genes, *RICTOR, MARCHF8*, and *BMP2*), while those in factors 2 and 3 were all positive. None of the genes associated with the CpGs (given by *Illumina 450k* platform) selected are in the set of mRNA features selected.

#### 4.1.2 Gene Set Enrichment Analysis (GSEA)

Given the overall directionality of the loadings of the features selected, GSEA was performed accordingly.

The results of the top-level Reactome pathways for the mRNA features selected are in Fig. 5.

**Figure 5:**
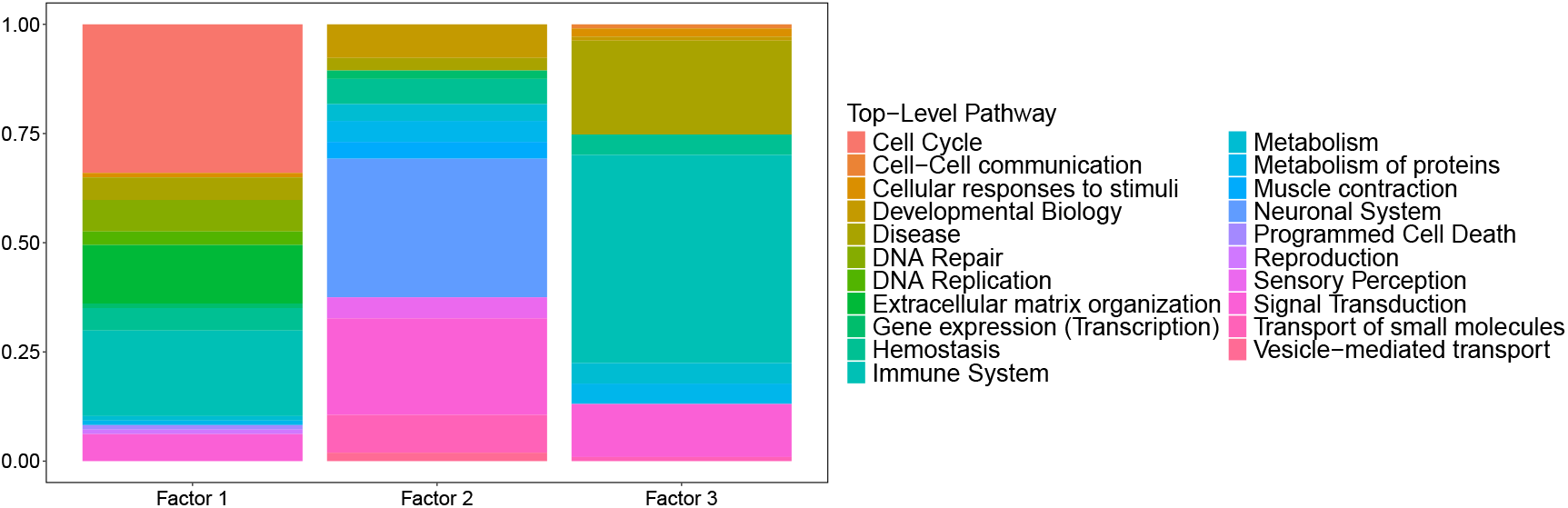
Proportion of top-level Reactome pathways identified by GSEA from ranked mRNA feature loadings across MOFA factors.

Overall, factor 1 captures a heterogeneous mix of biological processes, with enrichment in the immune system, cell cycle, and other functional categories like extracellular matrix organization, while factor 2 reveals a distinct functional profile, pointing toward a key role of the neuronal system. Also, signal transduction processes appear to play an important role in this factor. Factor 3 is specifically associated with immune-related functions. It also has a large proportion of pathways attributed to the ‘Disease’ top-level category, most of which correspond to cancer.

The top 15 enriched Gene Ontology biological processes associated with factors 1 and 3, based on the overall sign of their most important features, are shown in Fig. 6.

**Figure 6:**
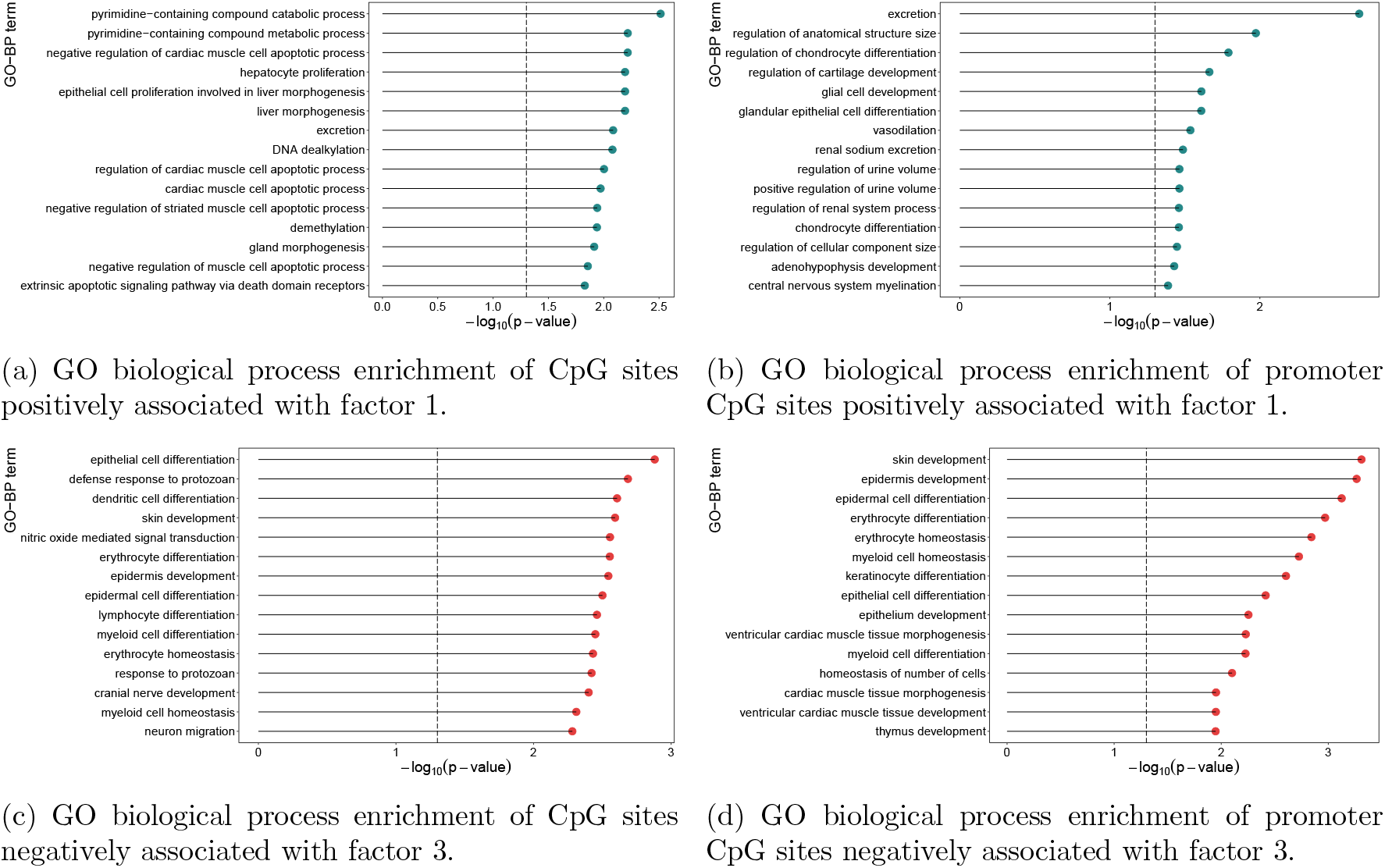
Gene ontology biological process (GO-BP) enrichment results for DNA methylation features associated with factors 1 and 3. Only pathways with more than 5 and fewer than 500 genes were tested. Significance was assessed at a threshold of 0.05.

The CpGs associated with factor 1 are predominantly enriched in pathways related to cell apoptosis, with additional involvement in metabolic and epigenetic processes. Focusing on promoter regions, the methylation changes in factor 1 CpGs are primarily associated with pathways involved in tissue development and cell differentiation, which is especially relevant given the glial origin of LGG (the ones with factor 1 positive). Additionally, several homeostasis-related pathways are present.

The pathways enriched by factor 3 highlight a strong theme around cell differentiation and immune system function. These include the development of various cell types, such as epithelial, epidermal, dendritic, erythrocyte, lymphocyte, and myeloid cells. A similar pattern is observed in promoter regions, and given that the same biological functions are enriched among factor 3 genes, this suggests that methylation at these CpG sites is likely associated with gene silencing.

### 4.2 Interpretation and Evaluation of the Findings

The CpGs along with the genes *RICTOR, MARCHF8*, and *BMP2*, selected by factor 1, were found to be upregulated in LGG samples. In contrast, the remaining 27 genes exhibited higher expression levels in GBM samples. Notably, increased CpG methylation and mRNA gene expression were significantly associated with improved survival.

The selected factor 3 genes are consistently upregulated in astrocytoma compared to oligodendroglioma samples, supporting the hypothesis that factor 3 is involved in the separation of the LGGs. An exception is observed for three genes — *BHLHE41, ADRB2*, and *ATP8B4* — which, although statistically significant, display log_2_ fold-change values slightly below the threshold of 1. Within the LGG group, all but five are significantly associated with survival. GBM samples do not show a consistent pattern in the features associated with factor 3, and, in fact, no gene shows significant differential expression between astrocytomas and GBM.

The genes associated with factor 2 seem to be linked to a less aggressive tumor phenotype. In fact, they are consistently downregulated when comparing astrocytomas and oligodendrogliomas to glioblastomas, suggesting a potential role in LGGs. However, no significant expression difference is observed between astrocytomas and oligodendrogliomas, likely because these genes are shared across specific subgroups within both tumor types. Indeed, a high expression of these genes within LGG is associated with better survival significance, except for three genes: *CAMKK1, DMTN*, and *STX1A*.

The genes and probes associated with the same factor appear to be connected, as evidenced by strong Pearson correlations among them and consistently low correlations with features from other factors. However, regarding gene regulation, the association of each probe with its gene is not that direct. From factor 3, eight probes were assigned to *ISM1* but do not seem to be regulating the gene, as evidenced in Fig. 7 for two of those probes.

**Figure 7:**
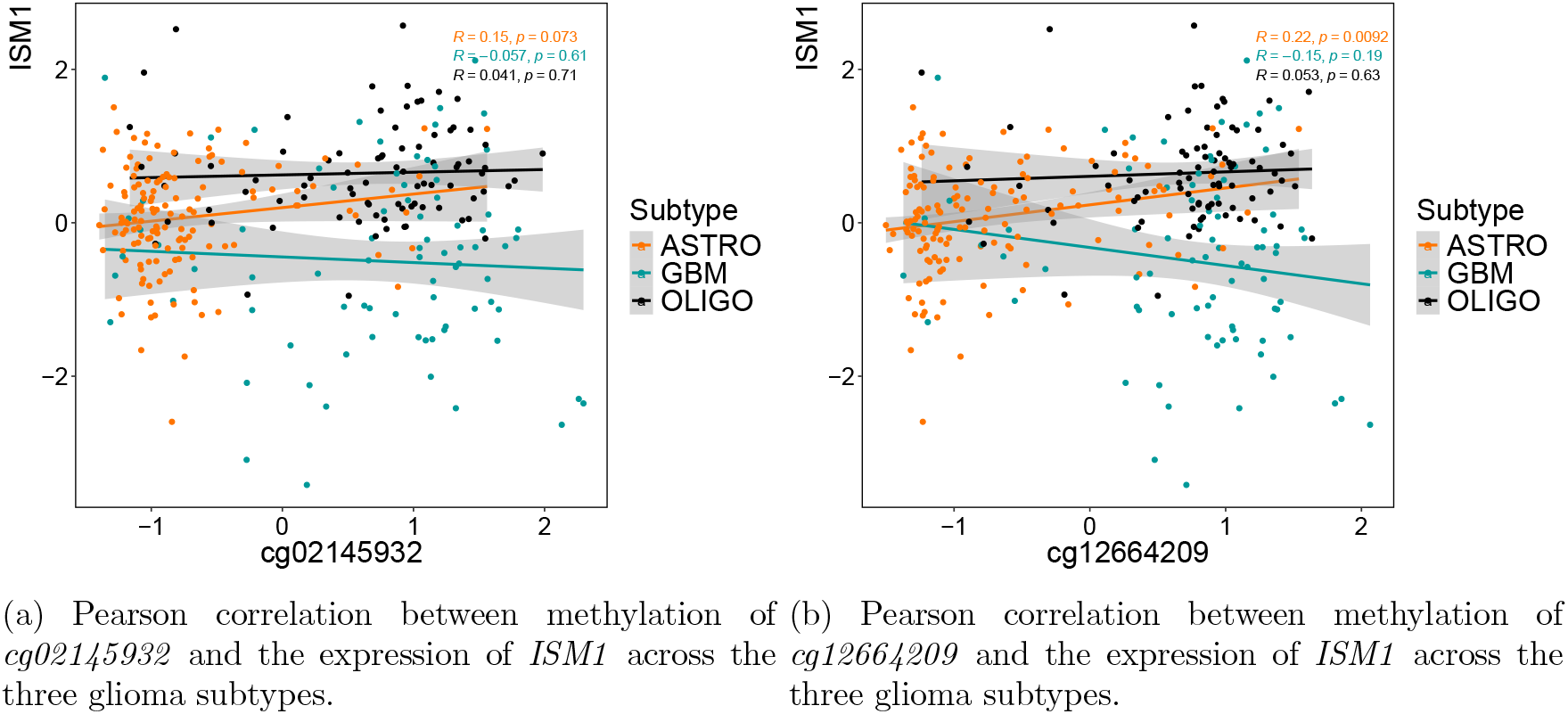
Correlation between methylation and gene expression for two probes annotated to *ISM1*. The data corresponds to the input matrices used in MOFA. Probe annotations are based on the Illumina 450K array.

Understanding how DNA methylation affects gene expression is highly complex [28, 29]. The effect of methylation is generally stronger when an entire region is methylated. Since eight probes that were selected target the same gene (*ISM1*), they may have a significant impact. By analyzing their correlation with the genes selected by factor 3, they show almost double the correlation, showing slightly stronger associations with the genes *SLC2A5, BLNK, TMEM119*, and *PLXDC2*. For the last gene mentioned, the correlation plots are in Fig. 8, with the same two probes given as an example above.

**Figure 8:**
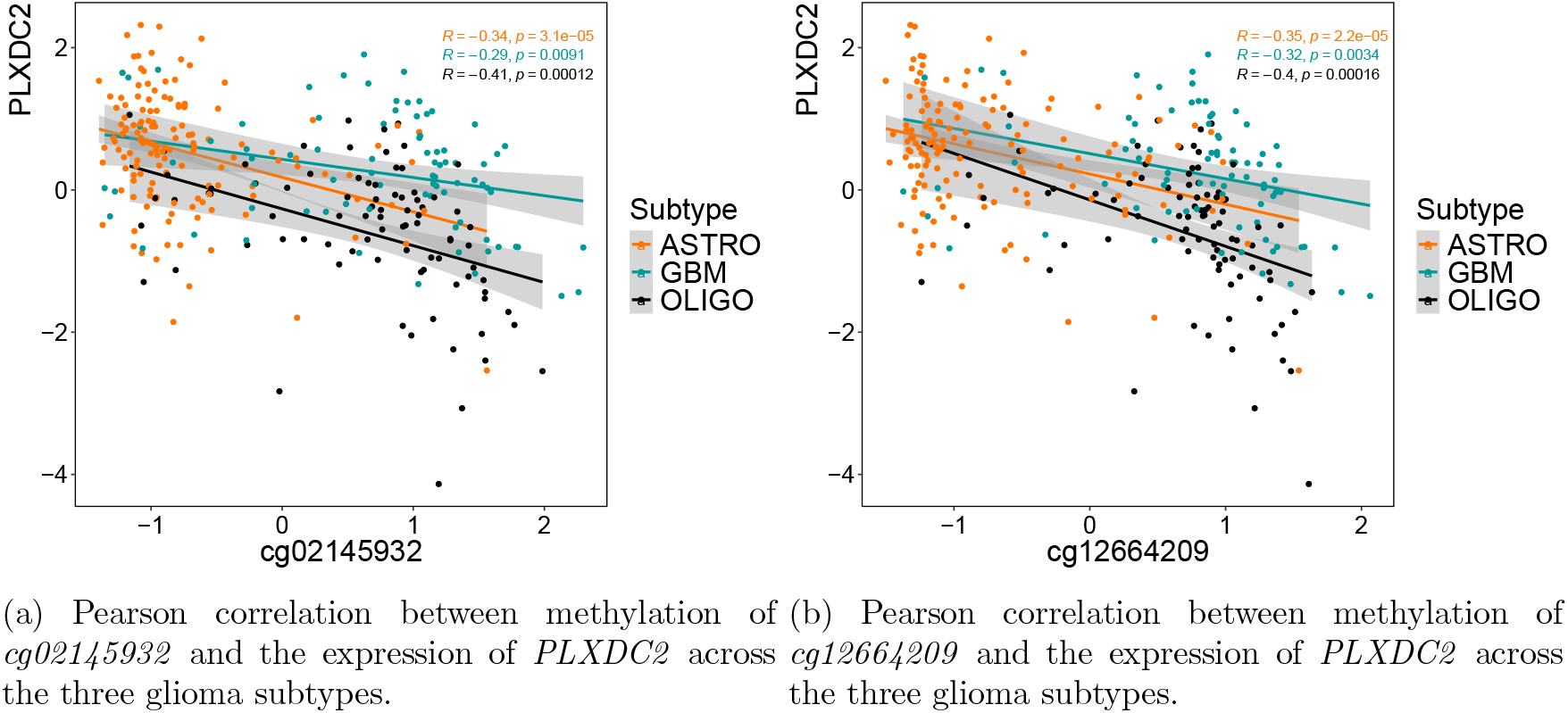
Correlation between methylation and gene expression for two probes annotated to *ISM1*. The gene *PLXDC2* was also selected by this factor. Correlations are computed using the input matrices to MOFA. Probe annotations are based on the Illumina 450K array.

### 4.3 Uncover MOFA-Derived Profiles

The MOFA factors appeared to be informative: factor 1 provided insights into the distinction between GBM and LGG groups, while factor 3 revealed heterogeneity specifically within the LGG group. Factor 2 was associated with high expression of neural system-related genes, suggesting an additional molecular profile beyond the three tumor labels. It was hypothesized that this profile might be linked to lower aggressiveness within the LGG group, prompting further investigation.

To explore this hypothesis, k-means clustering was applied using the three factors as input. The goal was to identify the number of clusters that best correlated with patient survival, specifically based on the p-values from the log-rank test. To account for variability due to k-means initialization, the analysis was repeated 30 times using 80% of the samples in each iteration, ensuring a balanced representation of censored and uncensored patients. The distribution of these p-values is shown in Fig. 9.

**Figure 9:**
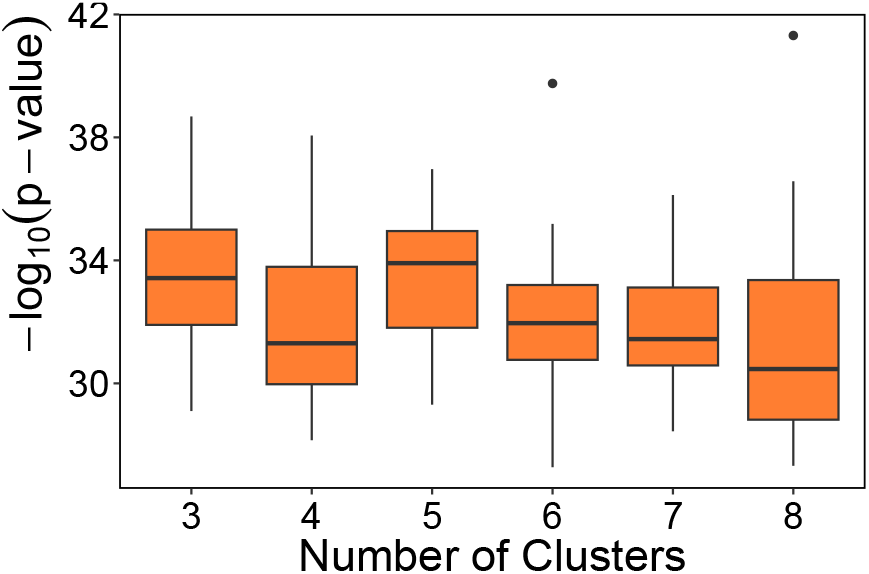
Distribution of the *−* log_10_(p-values) across different numbers of clusters tested.

The optimal number of clusters identified was five, although the difference in performance compared to using 3 clusters (the baseline) was not that much. The survival curves for three and five clusters are in Fig. 10 and the respective projections in MOFA factors 1 and 3 are in Fig. 11. The pairwise p-values of the five cluster solutions are in Table 2.

**Table 2:** Pairwise log-rank test p-values between clusters for the 5-cluster solution.

|  | Cluster 1 | Cluster 2 | Cluster 3 | Cluster 4 |
| --- | --- | --- | --- | --- |
| Cluster 2 | $8.98 \times 10^{-33}$ | — | — | — |
| Cluster 3 | $5.41 \times 10^{-1}$ | $3.85 \times 10^{-25}$ | — | — |
| Cluster 4 | $1.32 \times 10^{-11}$ | $5.41 \times 10^{-1}$ | $1.05 \times 10^{-9}$ | — |
| Cluster 5 | $2.66 \times 10^{-13}$ | $5.72 \times 10^{-1}$ | $7.12 \times 10^{-10}$ | $8.50 \times 10^{-1}$ |

**Figure 10:**
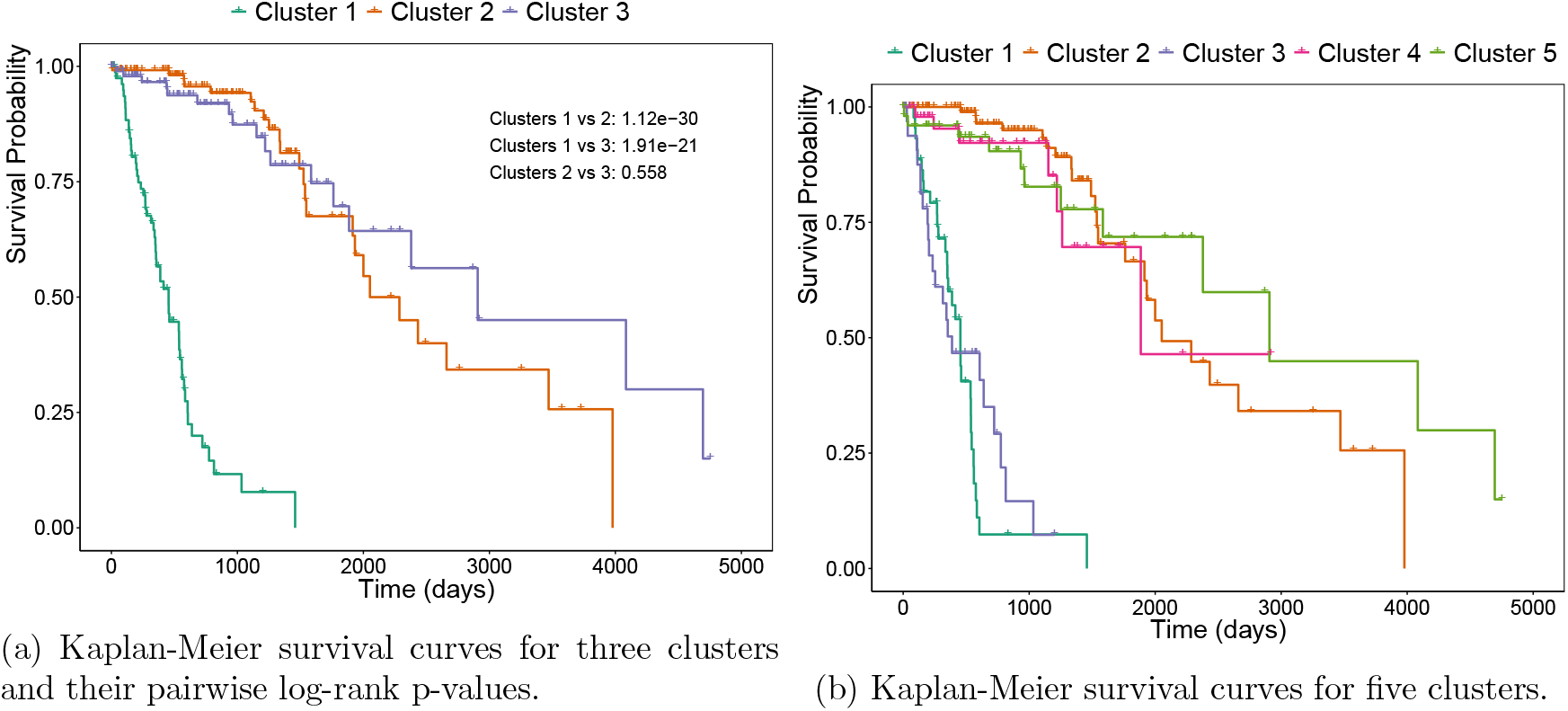
Kaplan–Meier survival curves for the 318 patients stratified by the three MOFA factors using k-means clustering.

**Figure 11:**
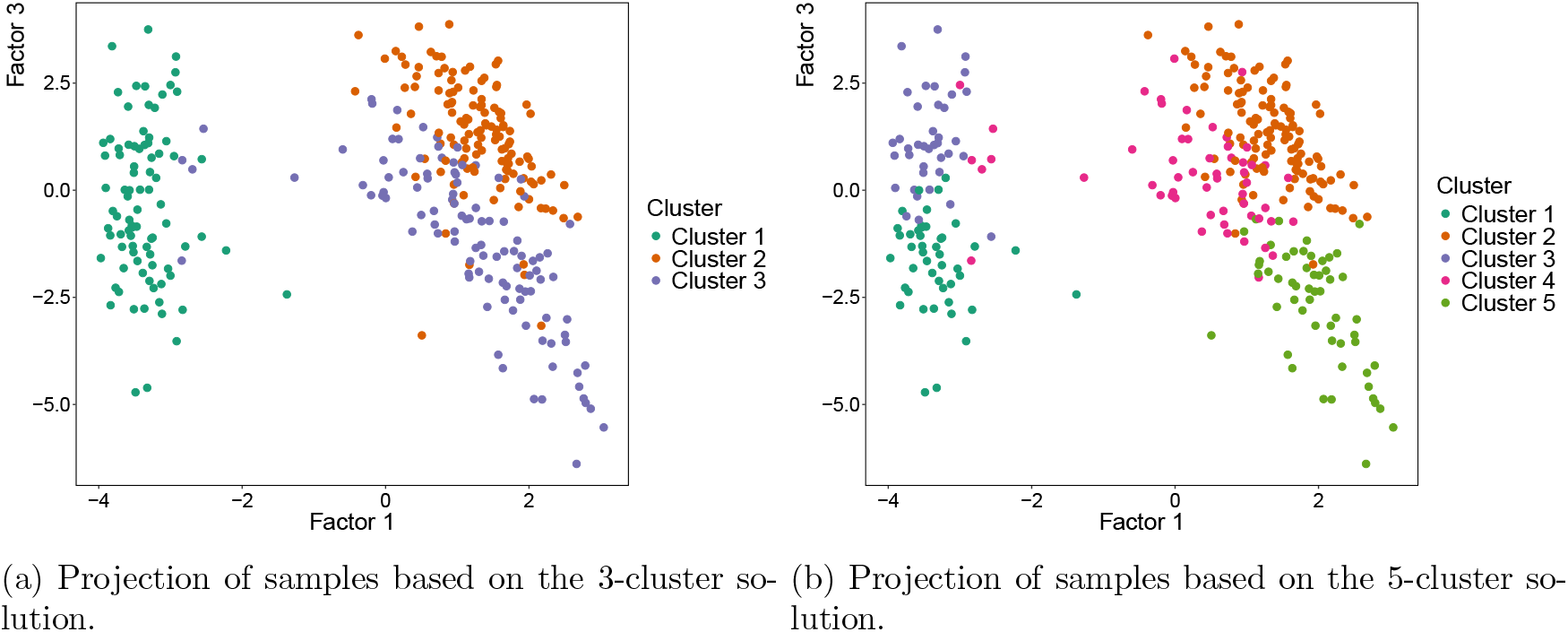
Sample projections of 318 patients according to cluster assignments with *k* = 3 and *k* = 5.

Table 3 summarizes sample distribution across the five clusters obtained with the described method, with proposed names GBM-1, ASTRO-2, GBM-3, MIX-LGG-4, and OLIGO-5. GBM cases belong mainly to clusters 1 and 3. Cluster 1 (GMB-1) contains the oldest patients and shows the highest percentage of deaths, possibly explaining its more aggressive survival profile compared to GBM patients in cluster 3 (GMB-3). This might be associated with a higher incidence of mutations in genes such as *EGFR*. Additionally, histological data reveal that most patients in cluster 1 are mostly diagnosed with primary GBM, whereas cluster 3 includes fewer primary GBM and more astrocytomas, further supporting the difference in clinical outcomes between these two GBM-enriched clusters. The majority of astrocytoma cases are grouped in cluster 2 (ASTRO-2), whereas oligodendroglioma cases were mainly found in cluster 5 (OLIGO-5). Cluster 4 (MIX-LGG-4) shows a balanced mix of LGG subtypes, suggesting an intermediate profile. Compared to the 3-cluster solution, the 5-cluster setting provides a clearer separation of LGG subtypes. In the 3-cluster case, one cluster grouped together 76 oligodendrogliomas and 25 astrocytomas, indicating limited resolution between these two subtypes. Notably, astrocytomas are often misclassified as GBM due to molecular similarities; however, this does not appear to occur with MOFA. In the 3-cluster solution, GBM samples were largely isolated into a single cluster, with only two samples assigned to the other two clusters.

**Table 3:** Summary statistics for each identified cluster, including clinical and molecular characteristics. For each cluster, we report the number of patients, mean age, proportion of males, and proportion of deceased individuals. The last columns display the distribution of glioma subtypes and the prevalence (%) of some key mutations.

|  | Number of<br>Patients | Age<br>(Mean) | Male<br>(%) | Dead<br>(%) | ASTRO | GBM | OLIGO | IDH1<br>(%) | PTEN<br>(%) | EGFR<br>(%) | TP53<br>(%) |
| --- | --- | --- | --- | --- | --- | --- | --- | --- | --- | --- | --- |
| GBM-1 | 45 | 62.87 | 55.56 | 71.11 | 0 | 44 | 0 | 0 | 33.33 | 31.11 | 17.78 |
| ASTRO-2 | 124 | 38.41 | 63.71 | 18.55 | 114 | 0 | 9 | 100 | 0 | 0 | 90.32 |
| GBM-3 | 39 | 55.89 | 42.11 | 56.41 | 0 | 34 | 0 | 0 | 30.77 | 23.08 | 17.95 |
| MIX-LGG-4 | 55 | 40.75 | 54.55 | 12.73 | 23 | 1 | 26 | 76.36 | 0 | 3.64 | 34.55 |
| OLIGO-5 | 55 | 48.11 | 49.09 | 23.64 | 5 | 1 | 49 | 89.09 | 0 | 0 | 10.91 |

DGE analysis was repeated using the mRNA and CpGs features selected by each MOFA factor and assessed within each cluster. The labels were changed to the respective group with the number of clusters at the end.

Factor 1 continue to capture features that separates high-grade gliomas from LGGs. Within GBM, factor 2 genes highlight strong differences between GBM-1 and GBM-3, with 24 Differential Expressed Genes (DEGs) between these two GBM groups. GBM-3 shares high expression of these genes with OLIGO-5 (0 DEG between them) and ASTRO-2 (1 DEG), suggesting that GBM-3 may exhibit a lower grade expression profile. Given the enrichment of these genes, GBM-3 could reflect a neural-like subtype of GBM, consistent with prior studies characterizing GBM subtypes [30]. The gene *SLC12A*, which exhibits the highest loading in this factor, has previously been associated with the neural subtype of GBM, reinforcing this interpretation [30]. Moreover, the MIX-LGG-4 cluster displays a distinctive and unique expression signature, showing 30 DEGs when compared with all other clusters, which potentially correspond to a transitional or mixed phenotype that does not align clearly with classic LGG or GBM categories. Factor 3 primarily differentiates the LGG subtypes, especially OLIGO-4 from ASTRO-2. OLIGO-5 exhibits a large number of DEGs (up to 30) and differentially methylated CpGs (up to 24) when compared to ASTRO-2 and GBM subtypes, indicating a well-defined epigenetic and transcriptional identity. Interestingly, GBM-3 shows greater molecular similarity to ASTRO-2 and a great dissimilarity with OLIGO-5 in this factor. On the other hand, GBM-1 appears to share some methylation and gene expression characteristics with MIX-LGG-4, suggesting potential epigenetic convergence between these groups.

## 5 Discussion

GBM tumors represent the most aggressive glioma subtype, exhibiting higher cell proliferation and invasiveness compared to LGG [9]. In this study, it was highlighted 27 genes overexpressed in GBM over LGG, predominantly linked to immune system activity, cell cycle regulation, and extracellular matrix organization - *FBXO17, RARRES2, JHY, PINLYP, AQP5, EVC2, XKR8, TOM1L1, EFEMP2, RBP1, CCDC8, FABP5, C9orf64, SH2D4A, CUL7, ARL9, CEP112, EID3, RAB34, RAB36, EMP3, SHROOM3, TSTD1, HDHD3, CMYA5, VASN* and *PRICKLE3*. Overexpression of *RAB34* is linked to poor prognosis, tumor invasion, and immune infiltration and is suggested as a possible immunotherapy target for glioma [31]. Though less studied, *RAB36* may share similar roles in vesicle trafficking, suggesting its potential relevance in GBM progression since they belong to the same RAB family of proteins.

The genes *RICTOR, MARCHF8*, and *BMP2* were found to be highly expressed in LGG tumors. All of them have both been reported in the literature as overexpressed in LGG, compared to GBM. The literature highlights the oncogenic role of the mTORC2 signaling pathway and its component *RICTOR* in GBM, with evidence that co-targeting *EGFR* and *RICTOR* leads to strong anti-tumor effects [32]. While *EGFR* alterations are known to activate the mTORC2 pathway, where *RICTOR* is a key component, this suggests that *RICTOR* overexpression in LGG may occur independently of *EGFR* mutation status, potentially reflecting alternative mechanisms of pathway regulation in LGG. In fact, this gene is in the Reactome pathway “Regulation of TP53 Expression and Degradation”, and its overexpression may contribute to enhanced degradation or reduced expression of *TP53*, a characteristic of the astrocytoma group. This set of 30 genes were identified as not only differentially expressed between the groups but also significantly associated with patient survival.

Three miRNA features were also found to be overexpressed in LGG: *MIR6071, MIR649*, and *MIR4666A*. Although they do not code for proteins, miRNAs play important regulatory roles in tumor biology, and their dysregulation impacts tumor suppressors and oncogenes [10]. *MIR6071* has been reported to be downregulated in GBM and is associated with tumor-suppressive functions, including the inhibition of cell proliferation, migration, and invasion. Notably, higher expression levels of *MIR6071* have been linked to better prognosis in glioma patients [33]. The roles of *MIR649* and *MIR4666A* are less well characterized; however, both are predominantly expressed in brain tissues, suggesting a potential relevance in glioma biology.

Additionally, DNA methylation patterns in LGG appeared particularly relevant to the regulation of cell apoptosis, especially at promoter regions enriched for genes involved in homeostatic processes and glial cell development. Although the precise regulatory impact of these methylation events is unknown, a strong correlation was observed between the methylation levels of the selected probes - *cg24041541, cg19070139, cg13727691, cg20826224, cg12630147, cg05163329, cg25604326, cg16590910, cg01764954, cg03974423, cg00428601, cg21174055, cg12604950, cg18276016, cg03609308, cg19113375, cg01818121, cg22230604, cg20525712, cg16483867, cg01338255, cg01843034, cg19998675, cg22451910, cg01118078, cg12907983, cg21940568, cg26448489, cg08409074* and *cg25251738* – and the expression of the 30 genes mentioned above: methylation was negatively correlated with genes overexpressed in GBM and positively correlated with the three LGG-specific genes. A higher methylation levels of these probes are associated with improved patient survival.

A molecular sub-profile within GBM was identified, characterized by high expression of neural-related genes and presented similarities to astrocytoma, regarding genetic expression and histological type. This less aggressive subgroup, found in younger patients, shows lower *EGFR* mutation rates. In fact, *EGFR* amplifications have been reported to define a much more aggressive tumor subpopulation, as it drives high-grade morphological features such as cell proliferation, invasion, and angiogenesis [10, 8]. This is because, as a potent oncogene, *EGFR* amplification activates the intracellular signaling pathways such as *PI3K/AKT/mTOR*, important regulators of cell growth and survival. Notably, *SLC12A5*, enriched in the cerebral cortex and linked to the neural GBM subtype [30], may play a vital role in the Central Nervous System. Other neuronal genes such as *ADAM11, RBFOX3*, and *HTR1E* may also indicate a favorable prognosis.

The same group of genes highlight a group within LGG, constituted by astrocytoma and oligodendroglioma patients. This group seems a transitional from the two groups, as 10 genes were found in a list of the top 100 genes upregulated in tumor with grade II compared to grade III, in *IDH1* mutated gliomas [34] - *ABLIM2, RYR2, ADAM11, RBFOX3, TMEM130, HTR1E, MPPED1, DMTN, MAP7D2* and *PACSIN1*. The other genes selected with them - *SLC12A5, DOC2A, NAPB, CPNE9, FADS6, MFSD4A, CAMKK1, SMIM10L2B, GFOD1, NEURL1, PHF24, CAMK2A, SNAP25, HIPK4, SULT4A1, PHYHIP, NCDN, DGKE, STX1A* and *CYP4X1* - are also upregulated in this group.

Astrocytoma and oligodendroglioma patients were well characterized. Astrocytomas showed high expression of immune-related genes, with 30 genes upregulated compared to oligodendrogliomas, 23 of which were significantly associated with survival (*SLC2A5, ADRB2, P2RY13, CSF2RA, SELPLG, IKZF1, ITGAM, LPAR5, SYK, RHBDF2, DOCK2, TMEM119, ATP8B4* and *TMEM52B, BHLHE41, LPCAT2, RGS10, RASAL3, ADAM28, IRF8, DEF6, WDFY4, PLCB2, APBB1IP* and *BLNK*). Interestingly, the less aggressive GBM subgroup shared this immune-like molecular profile with astrocytoma.

High methylation in oligodendroglioma affects genes linked to cell differentiation and immune function (for all regions and within promoter regions). Since similar genes are overexpressed in astrocytoma, this suggests methylation may silence them, indicating a possible interaction between these regulatory mechanisms. As noted in [29], promoter methylation is well known that is a key factor in gene silencing. In fact, several probes, eight out of thirty, were located near the gene *ISM1*, which is known for its role in anti-angiogenesis, immune response regulation, and the promotion of apoptosis—functions that highlight its anti-cancer potential [35]. However, methylation at these sites did not appear to directly regulate *ISM1*. Instead, a better correlation was observed with these immune-system function genes as *SLC2A5, BLNK, TMEM119*, and *PLXDC2*. However, these alterations may not be driven solely by these epigenetic modifications, as mutations could also play a contributory role.

Overall, the three glioma types were well distinguished, although astrocytoma can be challenging to classify due to overlapping genetic profiles with both GBM and oligodendrogliomas.

## 6 Conclusions

In this study, we integrated genomics, epigenomics, and transcriptomics data from TCGA to investigate glioma heterogeneity using multi-omics factor analysis. The analyses were based on the most recent WHO 2021 classification, which served as the reference standard for distinguishing astrocytoma, oligodendroglioma, and glioblastoma patients.

The identified factors combined information from the different omics layers, including genomics (mutations), epigenomics (DNA methylation), and transcriptomics (mRNA and miRNA), which allowed the reclassification of gliomas into five clusters, confirming numerous features previously associated with known groups while distinguishing between some glioma subtypes. For example, a distinct molecular profile within GBM, associated with the neural system and characterized by elevated expression of related genes, was identified, which may inform the development of novel prognostic markers and therapeutic strategies.

Supplementary analyses, including differential gene expression, gene set enrichment analysis, survival analysis, and Kaplan-Meier curves comparing the obtained groups, further corroborated the identified patterns at all the omics layers studied. Additionally, several novel features were identified that showed significant associations with survival.

The biomarkers identified in this study may help guide more personalized treatment strategies. Furthermore, considering the distinct molecular profiles observed within each glioma type, these biomarkers could prove especially valuable for improving disease classification, guiding treatment strategies, and enhancing prognostic assessments. Ultimately, the presented strategy allowed to generate testable hypotheses that can be explored in future studies to further validate their clinical relevance and applicability.

## Availability of data and materials

The R code developed for this analysis is open source and available at https://github.com-/CatarinaGameiroC/Article-MOFAinGliomas. The original datasets are not included due to their large size, but detailed instructions for downloading them are provided in the repository. The datasets are publicly available from The Cancer Genome Atlas (TCGA) database at https://portal.gdc.cancer.gov. The glioma classification using the WHO-2021 taxonomy guidelines is available at https://github.com/sysbiomed/MONET.

## CRediT authorship contribution statement

**Catarina Gameiro Carvalho:** Methodology, Software, Validation, Formal analysis, Investigation, Data Curation, Writing - Original Draft, Writing - Review & Editing, Visualization **Alexandra M. Carvalho:** Methodology, Formal analysis, Writing - Review & Editing, Supervision, Project administration **Susana Vinga:** Conceptualization, Methodology, Formal analysis, Writing - Review & Editing, Supervision, Project administration.

## Funding sources

Work supported by national funds through Fundação para a Ciência e a Tecnologia, I.P. (FCT) under projects UIDB/50008/2020 (IT), INESC-ID (UID/50021/2025 DOI: 10.54499/UID/50021/2025 and UID/PRR/50021/2025 DOI: 10.54499/UID/PRR/50021/2025), LAETA (DOI: 10.54499/UIDB-/50022/2020), SYNTHESIS (LISBOA2030-FEDER-00868200 - Projeto N° 15030) and TRACI (DOI: 10.54499/2023.17447.ICDT).

## Declaration of competing interest

The authors have declared no competing interests.

## Declaration of generative AI and AI-assisted technologies in the writing process

During the preparation of this work, the authors used ChatGPT to improve the readability and language of the manuscript. After using this tool/service, the authors reviewed and edited the content as needed and take full responsibility for the content of the published article.

## Acknowledgments

The authors thank Marta B. Lopes (FCT - NOVA), Bruno M. Costa (ICVS - UMinho) and Eduarda P. Martins (ICVS - UMinho) for their valuable biological insights to this work.

## Notes

https://github.com/CatarinaGameiroC/Article-MOFAinGliomas

